# Dynamic coordination of the perirhinal cortical neurons supports coherent representations between task epochs

**DOI:** 10.1101/514612

**Authors:** Tomoya Ohnuki, Yuma Osako, Hiroyuki Manabe, Yoshio Sakurai, Junya Hirokawa

## Abstract

Cortical neurons show distinct firing patterns across multiple task-epochs characterized by distinct computational aspects. Recent studies suggest that such distinct patterns underly dynamic population code achieving computational flexibility, whereas neurons in some cortical areas often show coherent firing patterns across epochs. To understand how such coherent single-neuron code contribute to dynamic population code, we analyzed neural responses in the perirhinal cortex (PRC) during cue and reward epochs of a two-alternative forced-choice task. We found that the PRC neurons often encoded the opposite choice-directions between those epochs. By using principal component analysis as population-level analysis, we identified neural subspaces associated with each epoch, which reflected coordinated patterns across the neurons. The cue and reward epochs shared neural dimensions where the choice directions were consistently discriminated. Interestingly, those dimensions were supported by dynamically changing contributions of individual neurons. These results indicated heterogeneity of coherent single-neuron responses in their contribution to population code.

## Introduction

Individual neurons across cortical areas show temporally flexible responses to multiple task-epochs characterized by different computational aspects, such as cue, action, and reward^1–3^. Recent studies have indicated that such diverse single-neuron responses are only interpretable in terms of their contribution to population dynamics which flexibly realize different computations, particularly in association and motor cortices^4–9^. These studies highlight the complex changes occurring in neural responses across epochs of a given task, which can provide orthogonal neural subspaces for independent computations. In contrast, individual neurons in many cortical areas have been shown to often encode relevant information across multiple epochs (Anterior cingulate cortex^10^, Insular cortex^11^, Motor cortex^12^, Orbitofrontal cortex^13–15^, and Perirhinal cortex^16^), suggesting their ability to support coherent representations through different task-epochs. Because of the differences in the forms of explanation in these studies (that is, population or single-neuron level), how coherent representations carried by individual neurons can be reconciled with dynamically changing population structure has not been well investigated.

In the present study, we explored the neural responses in the perirhinal cortex (PRC), which has been implicated in associative memory ^17–22^. This region receives sensory inputs from almost all modalities, reward-related signals from the amygdala and contextual information from the prefrontal cortex, entorhinal cortex and hippocampus^23–25^. Recent studies have shown that the PRC neurons modulate their sustained responses to visual cues as a function of time contexts^26–27^. It has also been shown that many neurons in the PRC change their tuning profiles between cue and choice-response epochs in a complex manner^14^. These results strongly suggest the capacity of the PRC neural population to employ both population dynamics and coherent representations through multiple task-epochs.

To investigate how the PRC shows coherent single-neuron code and dynamic population code across different epochs, we employed a standard two-alternative forced-choice task and analyzed neural responses in two epochs, where different computations are demanded: making predictions about the outcome of choices (cue epoch) and reinforcing the choices (reward epoch). By taking advantage of the interleaved visual and olfactory cue stimuli, which allowed us to evaluate modality-independent encodings, we analyzed dynamic population encodings related to different choices during those epochs in relation to single-neuron level selectivity.

## Results

Neurons in the PRC encode choice directions during a two-alternative forced-choice task. We trained rats to perform a two-alternative forced-choice task where they chose a target port (left/right) associated with a presented cue to obtain reward (Fig. 1a–b). The task performance was of a similar level regardless of the cue modality (mean correct rate in visual trials = 95.6 ± 5.5%; olfactory trials = 92.3 ± 4.3%). We recorded spiking activities from the left PRC (*n* = 207 neurons) during the task performance (37 sessions in five rats).

**Figure 1.**
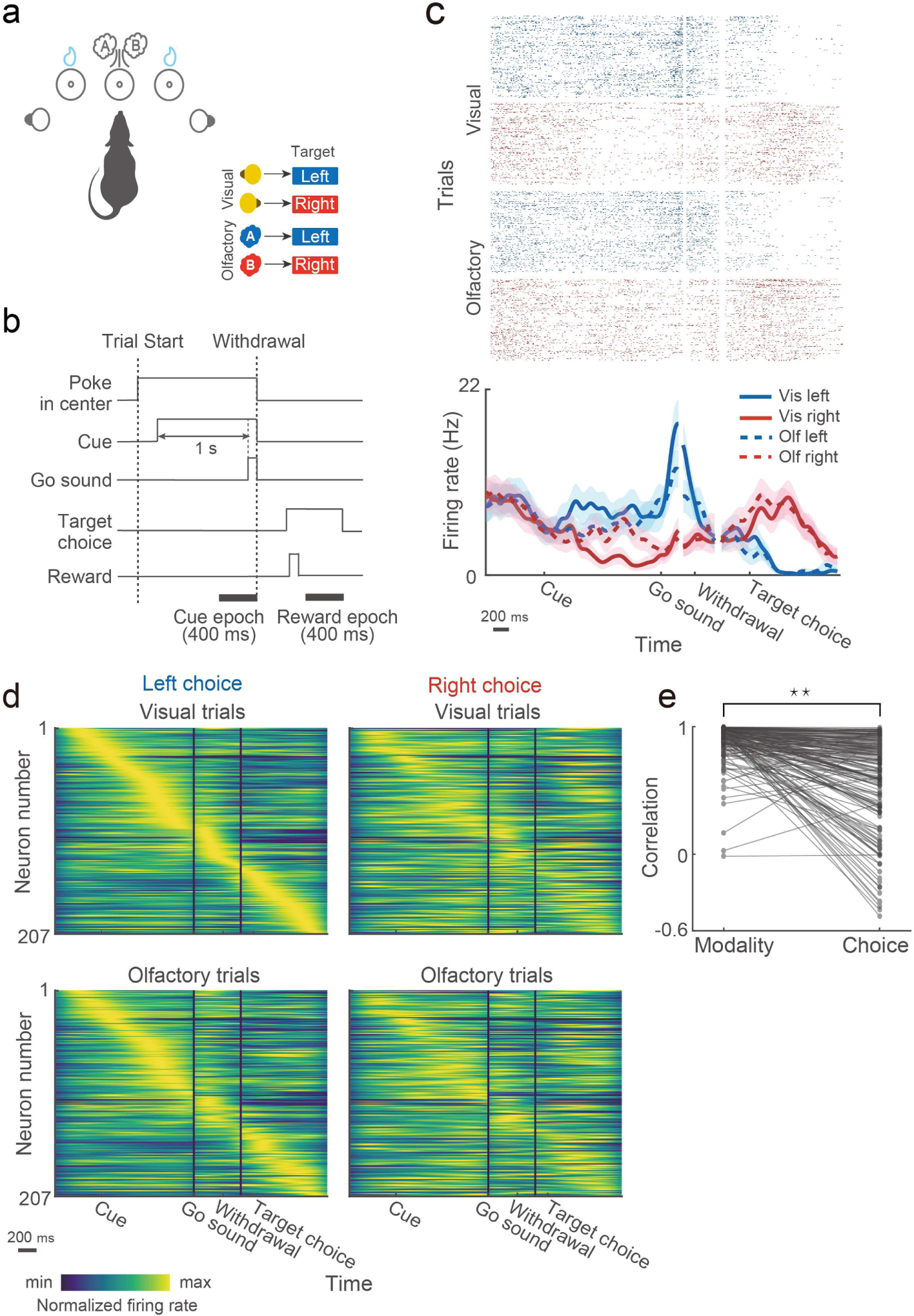
Firing patterns of the PRC neurons in a two-alternative forced-choice task. (a) Schematic drawing of the behavioral apparatus and the cue-target associative relationships. (b) Schematic of the task timeline. (c) Raster plot and peri-event time histogram showing response of a representative neuron. Trial types are classified according to cue modality and target choice as follows: blue, left target-choices; red, right target-choices; solid line, visual trials; dashed line; olfactory trials. Neural responses in correct trials were independently aligned to cue, withdrawal and target-choice onset and then reconstructed because of variable time between them. Lines and shaded areas indicate mean and s.e.m., respectively. (d) Firing patterns across all the neurons (*n* = 207) for the different trial conditions. In each trial type, mean firing rate of each neuron was normalized to its peak. For all trial types, the neurons were sorted by their peak firing time in visually-cued left choice trials (upper left). (e) Comparison of temporal firing patterns of individual neurons between the different modalities and choice directions. For comparison between different modalities, for each neuron, correlation coefficients were computed between peak-normalized firing rates in visual left (upper left in d) and olfactory left trials (bottom left in d). For comparison between different choices, correlation coefficients were computed between peak-normalized firing rates in visual left (upper left in d) and visual right trials (upper right in d). Nearly identical results were achieved when correlations were computed between the visual right and olfactory right trials and between the visual right and olfactory right trials (data not shown).

As shown in Fig. 1c, the PRC neurons typically showed distinct temporal firing patterns in left and right trials. To characterize how the PRC was activated by different trial conditions, we compared firing pattens among different cue-modalities and choices across all the recorded neurons. The neurons were sorted by their peak firing rates in visually-cued left choice trials (top left in Fig. 1d). As consistent with previous studies in other brain regions^28–33^, the peak responses of the PRC neurons tiled the duration of a trial. The response patterns across the neurons were well preserved between the cue modalities but much less so between the choice directions (comparison between the top and bottom in Fig. 1d). Conversely, the neurons frequently showed different response pattens between the choice directions (comparison between the left and right in Fig. 1d). We found that the majority of the neurons showed more strongly correlated response patterns between different modalities than between different choice directions (two-sided Wilcoxon signed-rank test; *P* = 1.1864×10^−24^; Fig. 1e). These results suggest that firings of the PRC neurons are more sensitive to the animals’ choice behavior than cue information.

We thus quantified selective responses of the individual neurons to the different choice-directions by using ROC analysis. We defined “choice-direction selectivity” broadly as signals reflecting a chosen direction in each trial. The individual PRC neurons encoded the choice directions at different time points in a trial (Fig. 2a for visual trials; Supplementary Fig.1 for olfactory trials). In addition, each neuron often showed such encodings at the time points other than its peak selectivity, suggesting that the individual neurons flexibly respond to multiple epochs of the task. We found two epochs where the choice-direction selectivity in both modalities reaches its peak (Fig. 2b), the cue epoch (−400 to 0 ms before withdrawal from the central port) and reward epoch (200 to 600 ms after choice). Many neurons encoded the choice-direction information during these epochs (Fig. 2c; 51.21% and 70.53% of the neurons showed significant selectivity in the cue and reward epochs, respectively). We observed a slight bias toward ipsilateral choice in the cue epoch (two-sided sign test; mean visual choice-direction selectivity = 0.006 ± 0.124, *P* = 0.8894; mean olfactory choice-direction selectivity = 0.0250 ± 0.102, *P* = 5.104×10^−4^) but no bias in the reward epoch (two-sided sign test; mean visual choice-direction selectivity = −0.026 ± 0.154, *P* = 0.2109; mean olfactory choice-direction selectivity = −0.021 ± 0.154, *P* = 0.3305). The choice-direction selectivity was highly consistent across the cue modalities during both epochs (*r* = 0.593, *P* = 5.1803×10^−21^ for the cue epoch; *r* = 0.858, *P* = 2.8367×10^−61^ for the reward epoch; Fig. 2c), and the PRC neurons were not tightly clustered as a choice-direction selective subpopulation and the others but rather showed graded selectivity as reported in other cortical areas^7, 34^. The neurons which showed significant selectivity across both modalities were sparser in the cue epoch (26% of selective neurons) than the reward epoch (48% of selective neurons), but we found that the neurons classified as non-selective during the cue epoch (48.79%, 101 of 207 neurons) showed moderate correlation between the cue modalities (*r* = 0.357, *P* = 2.5054×10^−4^; inset of Fig. 2c). This suggested even such neurons might convey a fraction of the choice-direction information. Altogether, these results suggest that sensory inputs from different modalities evoke similar response patterns across the PRC neurons according to learned cue-target associative relationships.

**Figure 2.**
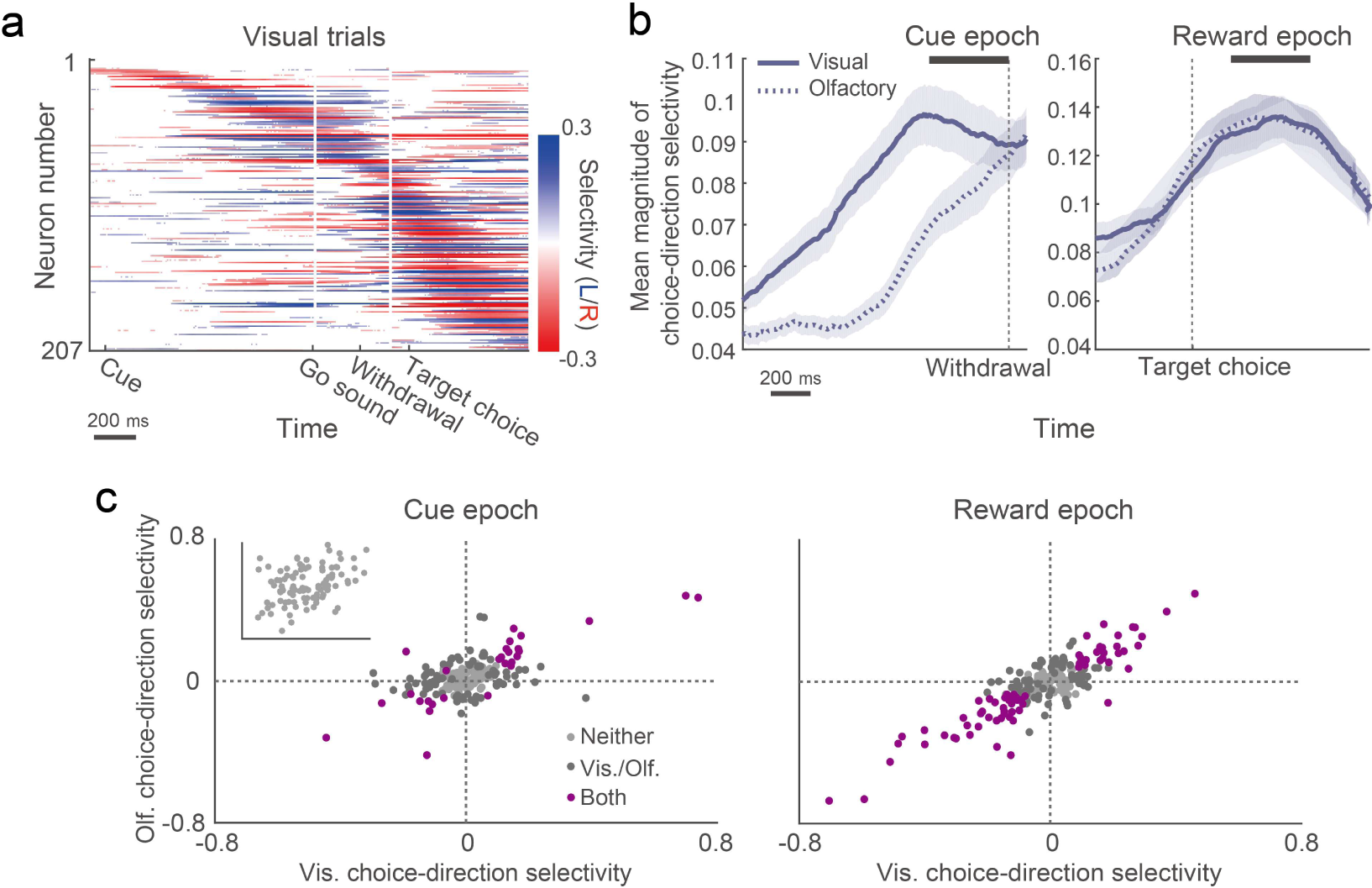
Choice-direction selectivity of the individual PRC neurons in different epochs of the task. (a) Temporal patterns of choice-direction selective responses of the PRC neurons (*n* = 207) in correct visual trials. Colors indicate selectivity to the left (blue) or right (red) choice. Neurons were sorted according to the time of their peak selectivity. Only segments with significant selectivity are shown (*P* < 0.05; 1,000 permutations). (b) Time course of the choice-direction selectivity magnitude in visual (solid line) and olfactory (dashed line) trials averaged across all the neurons. Neural responses around the withdrawal onset (left) and target-choice onset (right) are shown. Lines and shaded areas indicate mean and s.e.m., respectively. (c) Scatter plots showing the choice-direction selectivity of the neural population during the cue (left) and reward (right) epochs (*n* = 207 neurons). Each point corresponds to values of a single neuron. Colors indicate significance (*P* < 0.05; 1,000 permutations): light gray, no selectivity; deep gray, significant in either cue modality; purple significant in both cue modalities. Inset, the choice-direction selectivity of the non-selective neurons (*n* = 101) in the cue epoch.

A potential caveat in this conclusion, however, is that the choice-direction selectivity in the PRC can be attributed to some fundamental behavioral or contextual variables such as body posture, non-orienting movements, and spatial view^30, 35–39^. To evaluate the possible influence of those fundamental variables, we performed two control experiments. First, we monitored head angles of the animals (*n* = 2) during neural recordings (Supplementary Fig. 2a–c) using a head-mounted accelerometer^40^. To quantify the influence of body posture and view angle on the PRC neural responses (*n* = 105), we compared prediction performance between linear-regression models with choice and x-axis (interaural axis) head angle by computing correlation between the neural responses and the model prediction across trials (Supplementary Fig. 2d). The performance of the model with choice was higher than the model with x-axis head angle in both of the cue and reward epochs by 4.28% ± 10.98% (m.a.d) and by 32.07% ± 34.8% (m.a.d), respectively. We found that responses of the majority of the PRC neurons was better explained by the choice than the x-axis head angle (two-sided sign test; cue epoch, *P* = 0.0192; reward epoch, *P* = 1.3653×10^−7^). Second, a delay period was inserted between the choice onset and the reward onset to dissociate the influence of anticipatory licking behavior^41^ and spatial view (position) from the choice-direction selectivity in the reward epoch. We found that many of the neurons (69.52%, 73 of 105 neurons) encoded choice-direction after the onset of the reward (0 to 400 ms after reward onset) in accordance with the reward-epoch responses shown in Fig. 2c. The majority of these selective neurons (71.23%, 52 of 73 neurons) showed stronger selectivity after the onset of the reward than during the reward delay period (two-sided sign test; median difference in the magnitude of selectivity = 0.08, *P* = 3.7134×10^−4^; Supplementary Fig. 2e). Although the other neurons (28.77%; 21 of 73 neurons) showed stronger selectivity during the reward-delay period than after the reward onset, the size of such bias (that is, difference in magnitude between these periods) was relatively small (0.058 ± 0.0609; *n* = 21 neurons) as compared to the bias to the reward onset (0.1586 ± 0.1217; *n* = 52 neurons). Taken together, these results suggest that fundamental variables such as body posture, non-orienting movements, and spatial view themselves are not major factors for the choice-direction encodings in the PRC.

Dynamic temporal encoding patterns in the PRC. Thus far, we showed that the PRC neurons encoded the choice directions across the different cue modalities and that such signals were apparent at multiple epochs of the task (Fig. 2 and Supplementary Fig.2). Given the reduced correlation of temporal response patterns between the different choices (Fig. 1d–e), it is possible that the individual neurons flexibly tuned to different choices at different time-points of the trial duration. To characterize how each neuron represented the choice directions over the trial duration, we classified the neurons into two groups based on their peak selectivity: left-selective and right-selective neurons. As shown in Fig. 3a, we found similar numbers of selective neurons for both choice-directions. As expected, individual neurons showed selectivity to the opposite choice direction in time points other than their peak responses. Remarkably, when we sorted these neurons by their peak responses to the opposite choice (that is, left-selective neurons by right peaks and right-selective neurons by left peaks), response patterns nearly tiling the entire trial duration appeared (Fig. 3b). These results suggested flexible engagements of individual neurons for different choice-directions rather than recruitment of distinct subpopulations for each choice-direction.

**Figure 3.**
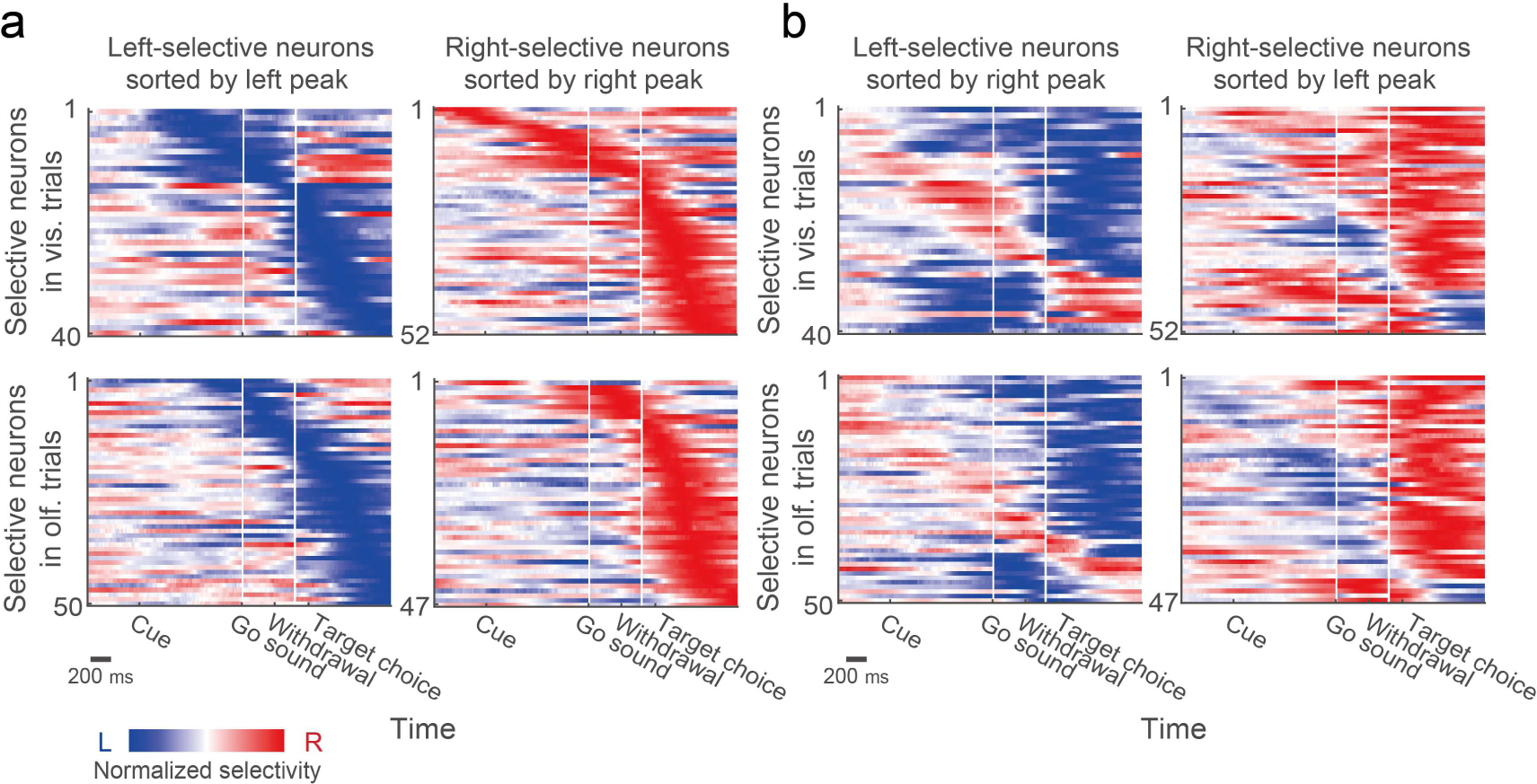
Time-varying choice-direction selectivity in individual neurons. (a) Temporal response patterns of left-selective and right-selective subpopulations. In each modality, neurons were classified by their peaks of significant choice-direction selectivity (Fig. 2a and Supplementary Fig.1). The selectivity of each neuron was normalized to its peak. The neurons were sorted according to the time of their peak selectivity. To visualize the entirety of time-varying selectivity, all segments with and without statistical significance are shown. (b) Same as in a, except for that the neurons were sorted by their peak selectivity to the opposite choice direction.

We next sought to understand how such dynamic neural encoding patterns in the PRC appear and evolve through the trial duration as a population. A time-resolved pattern analysis^42^ (Methods) was performed to visualize temporal evolution of the choice-direction encoding patterns across the PRC neurons (*n* = 207). Given the overall correlation of the choice-direction selectivity between the cue-modalities during the epochs where such encodings peak (Fig. 2 b–c), we focused on modality-independent encoding patterns (for comparison between the visual and olfactory trials, see Supplementary Fig. 3). The population response pattern evolved with two time-stable states (Fig. 4a). An encoding pattern was sustained during the presentation of the cue and was followed by a transient pattern during the movements towards the target ports. Soon after the rats chose a target port, the encoding pattern settled again into a stable state. To test for reliability of such encodings, we computed the mean performance of the classifier during the cue and reward epochs and compared them with a baseline epoch (−400 to 0 ms before the cue onset). As shown in Fig. 4b, the choice directions were decoded during both epochs above chance level (cue epoch: *P* ≈ 0.002; reward epoch: *P* < 0.001). Importantly, the encoding patterns were substantially inverted between the cue and reward epochs (scatter plots in Fig. 4a). The mean classification performance across those epochs revealed reliably inverted encoding patterns (*P* < 0.001; Fig. 4c). It is noteworthy that the inverted state was time-stable (Fig. 4a). This indicates that the time-varying encodings of the choice directions which we found in the previous section (Fig. 3b) are explicitly aligned to between the cue and reward epochs. In other word, the inverted encodings might be a form of coherent representations across different epochs of a given task, which is mediated by the individual PRC neurons.

**Figure 4.**
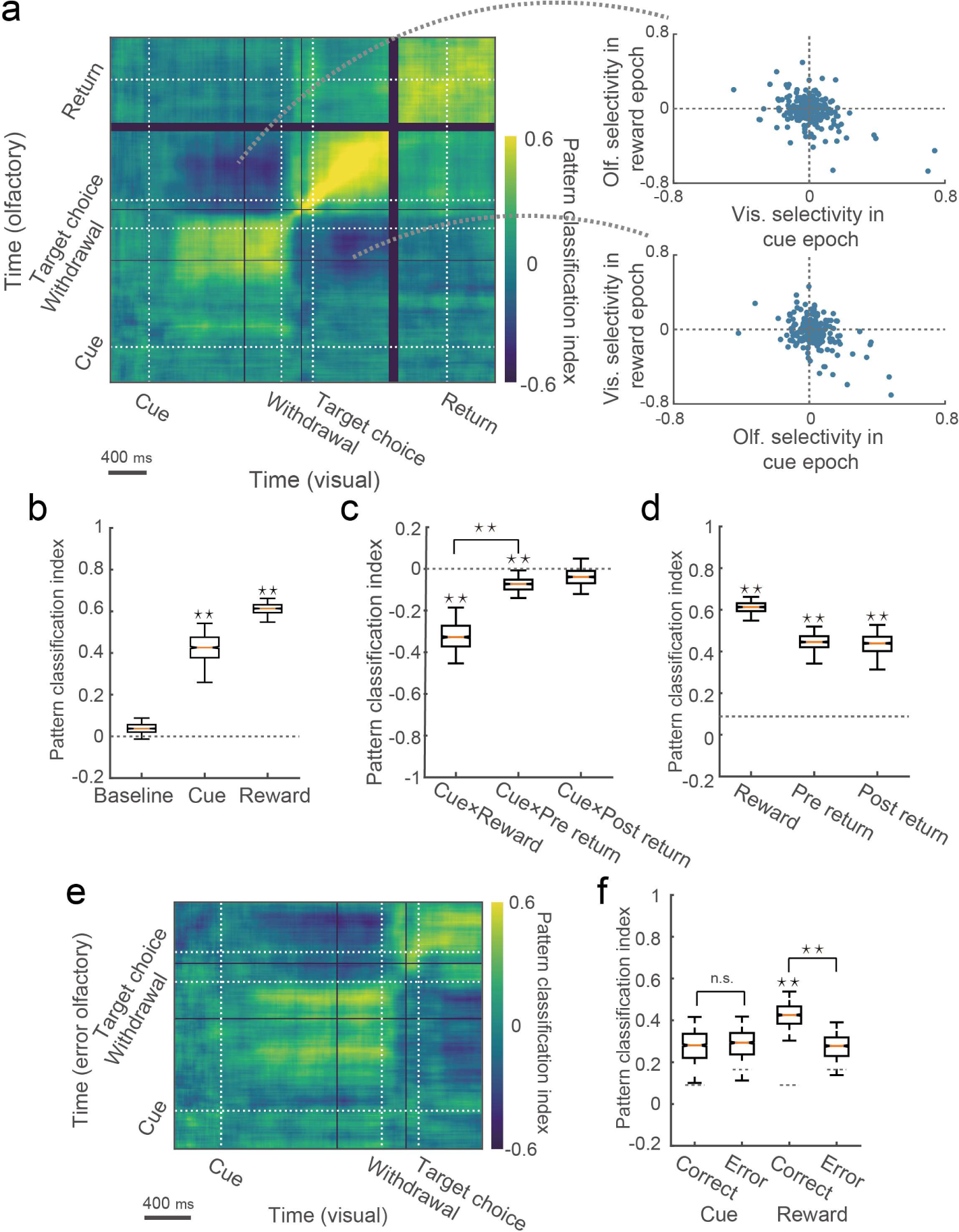
Time-resolved pattern analysis for temporal changes of choice-direction selectivity. (a) Performance of the pattern classification analysis (left). White lines correspond to the onset time of cue, withdrawal, target choice, and return movement to the central port (target-choice offset). We trained a classifier using neural responses in correct visual trials to discriminate choice directions and tested them with neural responses in correct olfactory trials (*n* = 207 neurons). Temporal inversions of choice-direction selectivity between the cue and reward epochs are shown by scatter plots (right). (b) Mean classification performance during the baseline, cue and reward epochs. (c) Mean classification performance across two different epochs. (d) Mean classification performance during the reward, pre-return and post-return epochs. Dashed line indicates the 97.5th percentile values of the baseline distribution shown in b. (e) A classifier tested with erroneous olfactory trials (*n* = 119 neurons). (f) Mean classification performance in erroneous trials compared with correct trials. Dashed lines indicate the 97.5th percentile values of the baseline distributions in the correct and erroneous trials. In box plots: orange line, median; box limits, 25th and 75th quartiles; notch limits, (1.57×interquartile range)/√*n*; whiskers, 95th percentile range (two-sided) of the distribution. Asterisks indicate statistical significance based on estimated *P* values (*P* < 0.05; Methods), and n.s. indicate insignificance.

However, it is still possible that the inverted encoding patterns were due to the reversal of orienting movements between the cue and reward epochs. To directly test this, we analyzed neural responses around the time when the animals left the target ports in preparation to the next trials (Supplementary Fig. 4): the pre-return (−400 to 0 ms before the choice offset) and post-return epochs (0 to 400 ms after the choice offset). These epochs did not show the equivalent level of inverted patterns from the cue epoch (cue × pre-return epochs: *P* ≈ 0.016; cue × post-return epochs: *P* ≈ 0.1858; comparison between cue×reward epochs and cue×pre-return epochs: *P* ≈ 0.023; Fig. 4c) despite the fact that the choice-direction information was robustly represented during both pre-return and post-return epochs (*P* < 0.001; Fig. 4d). These results indicated little influence of orienting movements on the dynamic choice-direction encoding patterns.

We also asked whether the behavioral performance affect neural responses by performing the pattern classification analysis with correct visual and erroneous olfactory trials (*n* = 119 neurons with sufficient number of erroneous trials; Fig. 4e). As shown in Fig. 4f the mean classification performance during both epochs was chance level in erroneous trials (error cue: *P* ≈ 0.1389; error reward: *P* ≈ 0.1349). Meanwhile, in correct trials, the mean classification performance was above chance for the reward epoch and marginally significant for the cue epoch (correct cue: *P* ≈ 0.0729; correct reward: *P* < 0.001). We tested whether the classification performance during each epoch decreased from the correct trials to erroneous trials and found significant reduction only in the reward epoch (cue: *P* ≈ 0.5385; reward: *P* < 0.001). These results indicate dynamic recruitment of the PRC neurons during the different epochs, which might flexibly support different computational demands in each epoch.

Analysis for population encoding structure in relation to single-neuron representations. To investigate how such coherent response patterns appeared and worked under the entire population structure, we identified neural subspaces for the cue and reward epochs by performing principal component analysis (PCA). Since our focus was to compare population response patterns between the different epochs, PCA was applied to time-averaged cue-epoch and reward-epoch responses across conditions (that is, PCA on neurons × conditions matrix for each epoch) ^43–44^. This allowed us to clarify the population-encoding structure derived from the cue-epoch responses and reward-epoch responses and directly investigate interrelationship between them. We projected the cue-epoch and reward-epoch responses onto the first two dimensions of the cue-epoch and reward-epoch subspaces (Fig. 5a and Supplementary Fig. 5a). The different choices and cue modalities were clearly separated when we projected the neural responses onto the corresponding neural subspaces (Fig. 5a, cue epoch in upper left; reward epoch in bottom right). Although we observed highly correlated choice-direction selectivity between the visual and olfactory cues (Fig. 2c), modality information was evident in these plots. We quantified separation among the four different conditions in those plots by comparing the average distance among responses under different conditions (across-condition distance) with the average distance among responses under the same conditions (within-condition distance). The former distance indicates discriminability of the different conditions and the latter indicates variability of the population responses within each condition. The results revealed that the across-condition distance was significantly larger than within-condition distance in both projections, indicating reliable encodings of the different conditions during both epochs (*P* < 0.001; Fig. 5b and Methods). We also determined whether these neural subspaces depended on the entire neural population or on only a fraction of the neurons. We found that neural weights were highly distributed across the neurons (Fig. 5d and Supplementary Fig. 5b) with no neurons showing zero weight. This suggest that these subspaces reliably reflect coordinated response patterns across the PRC neurons. To directly investigate the relationship between the cue-epoch and reward-epoch responses, we projected the population responses onto interchanged neural subspaces (that is, the cue-epoch responses onto the reward-epoch subspace and the reward-epoch responses onto the cue-epoch subspace)^4, 5, 8^. The results showed that the choice directions but not the cue modalities were separable through the epochs (Fig. 5a, cue-epoch responses onto reward-epoch subspace in bottom left; reward-epoch responses onto cue-epoch subspace in upper right). For those interchanged projections, we also quantified separations among different conditions by the same method as the corresponding projections. As shown in Fig. 5c, in both subspaces, we found cluster separations above chance (reward-epoch responses in cue-epoch subspace: *P* < 0.001; cue-epoch responses in reward-epoch subspace: *P* ≈ 0.003), revealing that the choice-direction information was reliably sustained between those different epochs. We also projected the neural responses during the baseline epoch onto the same neural subspaces as negative controls (Supplementary Fig. 5c). This analysis revealed no significant cluster separations (*P* ≈ 1 for cue-epoch subspace; *P* ≈ 1 for reward-epoch subspace; Fig. 5c), indicating no reliable encodings of the task information. We then tested whether the temporal response patterns of the individual neurons contributed to the observed discriminability by projecting randomly shuffled data where the temporal response patterns of the individual neurons were collapsed (Supplementary Fig. 5d). The results showed no significant separations (*P* ≈ 1 for shuffled reward-epoch responses in cue-epoch subspace; *P* ≈ 0.99 for shuffled cue-epoch responses in reward-epoch subspace; Fig. 5c), suggesting that the coherence between the cue and reward epochs at the single-neuron level was critical for the discriminability observed at the population level.

**Figure 5.**
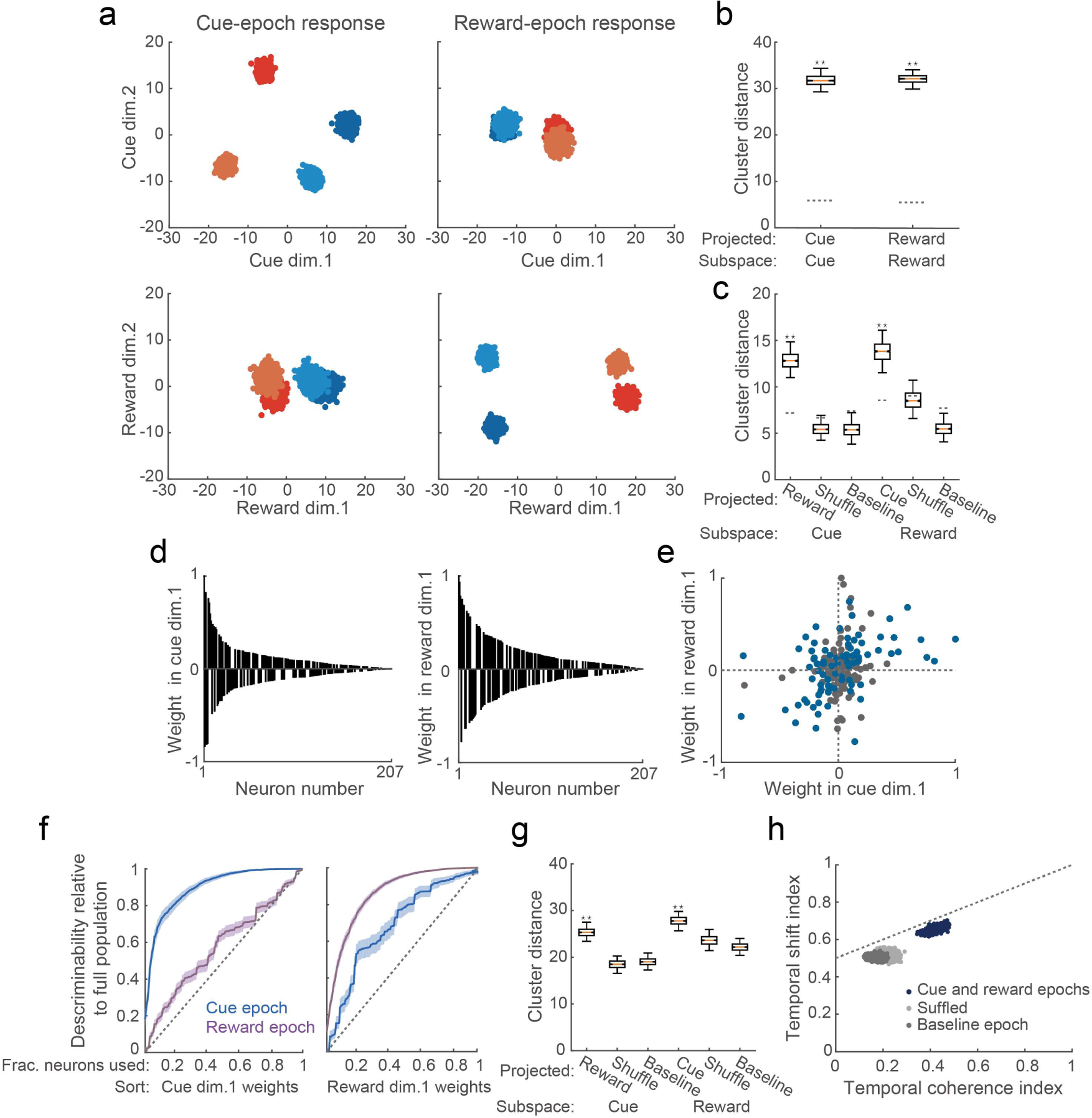
Principal component analysis of neural population responses. (a) Population responses during cue and reward epochs projected onto the first two dimensions of cue-epoch and reward-epoch subspaces. Blue, left target-choice in visual trials; pale blue, left target-choice in olfactory trials; red, right target-choice in visual trials; orange, right target-choice in olfactory trials. Each point corresponds to the population response in a subset of trials. (b–c) Comparisons between across-condition distances and within-condition distances. Dashed lines indicate the 97.5th percentile ranges of within-condition distances.(b) Distances obtained from projections onto corresponding subspaces. (c) Distances obtained from projections onto interchanged subspaces. (d) Weights of individual neurons in the first dimensions of the cue-epoch (left) and reward-epoch (right) subspaces shown in a. The values were normalized by magnitude of the highest weighting value and were ordered by magnitude. (e) Correlation between the neural weights in the first dimensions of the cue and reward subspaces shown in d. Blue points, neurons with significant choice-direction selectivity (*P* < 0.05) across the cue and reward epochs. (f) Choice-direction discriminability by neural populations with increasing numbers of neurons. Neurons were arranged by the weighting value on the first dimension of the cue epoch (left) or the reward epoch (right). Values were normalized to the discriminability by the full population. Lines and shaded areas represent mean and 95th percentile range, respectively. Dotted lines indicate increase expected by random incorporation of the neurons. (g) Temporal shifts in representational geometry. The same shuffled data as in d was shown here. We considered the baseline-epoch responses to be chance level. (h) Relationship between temporal coherence and dynamics of the PRC population responses (Methods). Vertical axis indicates proportion of the task representations sustained between different epochs, and horizontal axis indicates the degree of temporal shifts. Each single point corresponds to the population response in a subset of trials. The discriminability by the cue-epoch and reward-epoch responses is shown as blue points. The same for the baseline-epoch responses and shuffled responses are shown as deep gray and light gray points, respectively. In box plots, the same convention as in Fig. 4.

It might be possible that only a handful of neurons had strongly contributed to both cue-epoch and reward-epoch subspaces, thus resulting in the shared neural dimensions (that is, the first dimensions which consistently discriminated the choice directions) shown in Fig. 5a. However, this was not the case. As shown in Fig. 5e, the plot of the values of neural weights in the first dimensions of the cue-epoch and reward-epoch subspaces showed neither explicit clusters nor a tight correlation but rather exhibited a continuous distribution with a moderate correlation (*r* = 0.33, *P* = 1.51×10^−6^). This suggests that many neurons dynamically changed their contribution to the shared dimensions across the epochs, while a degree of temporal coherence supporting those shared dimensions were maintained at the population level. Diverse relationships of neural weights of the cue and reward epochs were observed even among neurons with significant choice-direction selectivity (computed using ROC analysis in Fig. 2c) across the epochs (blue points in Fig. 5e). This result indicated heterogeneity of temporally coherent single-neuron responses in their contribution to the entire population structure. In other words, each of those neurons might play unique role in population encodings under different computations.

To further establish population-level encoding^7, 34, 45–46^ in the PRC, we directly examined distribution of choice-direction information over individual neurons. We asked what fraction of the neurons is necessary to achieve choice-direction discriminability (that is, across-condition distance for different choice directions on the first dimension of a given epoch) which is equivalent to the entire population (Methods). The number of neurons incorporated were gradually increased, beginning with ones with the highest magnitude of the weighting values on the first dimension of the cue or reward epoch (Fig. 5f). As expected from the shape of the distribution in Fig. 5d, when the neurons were sorted by the cue-epoch weights, the choice-direction discriminability in the cue subspace steeply increased with the incorporation of neurons (left in Fig. 5f). However, the top 76.33% of the neurons was necessary to achieve the discriminability as high as the entire population. Similarly, when the neurons were sorted by the reward-epoch weights, the discriminability in the reward subspace reached as high as the full population after including 87.44% of the neurons (right in Fig. 5f). These results support population-level representations in which information is distributed across a number of individual neurons in the PRC. On the other hand, the choice-direction discriminability more gradually increased for the epochs to which the orders of neurons were not aligned (Fig. 5f), highlighting the dynamic changes in the neural weights between the epochs observed in Fig. 5e. We found that nearly all the neurons were necessary to achieve the discriminability as high as the full population (neurons sorted by cue-epoch weights, 100%; neurons sorted by reward-epoch weights, 98.55%), indicating that the choice-direction representations across the two epochs were distributed widely over the entire population. However, the mean discriminability in the epochs in which the neurons were indirectly sorted more steeply increased than the linear increase which is expected when the neurons were added in a random order (Kolmogorov-Smirnov test; cue epoch, *P* = 2.4985×10^−11^; reward epoch *P* = 0.0045). As shown in Fig. 5f, by incorporating a small subset of the neurons (∼10%), those curves deviated from the linear increase and the deviation was sustained through the almost all the parts of the curves. Therefore, the choice-direction information in the cue and reward epochs was highly distributed across the neurons, but its coherence was reliably sustained at the population level.

Efficient encoding of choice directions and different task-epochs by the PRC population. As shown in Fig. 5a, the population response patterns representing the choice directions were substantially inverted when we compared the cue-epoch responses and reward-epoch responses in the same neural subspace (comparison between horizontally arranged plots in Fig. 5a). In contrast, when the population responses from one of these epochs were projected onto the cue-epoch and reward-epoch subspaces (comparison between vertically arranged plots in Fig. 5a), the relative positions of the different choice-directions were preserved, indicating that the directions (that is, signs) of the first dimensions were consistent. Therefore, these results suggested that information of the different epochs was also represented in the shared dimensions by flexible changes in the representational geometry. The inverted geometry was consistent with the dominant temporal response pattern observed in Fig. 4a. To test reliability of the temporal inversions, in each subspace, we computed the average distance between population responses under the same conditions in the different epochs (for example, distance from cue-epoch responses to reward-epoch responses in the cue subspace) (Methods). We compared those distances with average distance from one of those epochs to the baseline epoch (for example, distance from cue-epoch responses to baseline-epoch responses in the cue subspace), which is equivalent to chance-level cluster shifts caused by the absence of task representations (that is, overlapped clusters at near zero in Supplementary Fig. 5c). Therefore, if the representational patterns are reliably inverted between the cue and reward epochs, then the temporal shift distances between those epochs should be significantly larger than the chance-level shift distances. As shown in Fig. 5g, the results showed larger task-epoch dependent shifts of the representational patterns than expected by chance (*P* ≈ 0.002 for cue-epoch subspace; *P* ≈ 0.009 for reward-epoch subspace). The discriminability was diminished when the temporal patterns of individual neural responses were collapsed by data shuffling (*P* ≈ 1 for shuffled reward-epoch responses in cue-epoch subspace; *P* ≈ 0.9021 for shuffled cue-epoch responses in reward-epoch subspace). These results suggested that the coherent response patterns at the individual neurons mediated flexible changes for task-epoch representations. We next asked how these flexible encodings for the cue and reward epochs (Fig. 5g) were related to the coherent choice-direction encodings across these epochs (Fig. 5c). We projected the PRC population responses onto the two axes capturing variances of those variables (temporal coherence index and temporal shift index in Fig. 5h; Methods). We found that variability of the population responses across subsets of trials were clearly aligned to the line that maximized both indices (dashed line in Fig. 5h), suggesting that the PRC population specifically tuned to support both coherence and dynamics of neural representation between the epochs. Transition of the population response between the cue and reward epoch resulted in a high correlation between the two indices (*r* = 0.629, *P* = 2.8077×10^−111^), but the correlation was absent in transition between the baseline epoch and the cue or reward epochs (*r* = −0.018, *P* = 0.5699).

Importantly, the correlation depended on the temporal response patterns in the individual neurons (*r* = −0.088, *P* = 0.0055 for shuffled responses), suggesting that coherent single-neuron encodings were critical to efficiently enhance coherence and dynamics of the neural representation. Taken together, these findings indicated that the PRC population reconciles coherent representations of choice directions with dynamic representations for different epochs via temporally flexible but structured neural responses (that is, inverted selectivity between epochs).

## Discussion

We investigated how coherent single-neuron representations across different epochs can be reconciled with temporal dynamics of population structure. Individual neurons in the PRC modulated their firings according to choice directions in each trial, and many neurons inverted such selectivity between the cue and reward epochs. Despite the structured temporal response patterns, we found dynamic reorganization of the population encoding structure between the different epochs (that is, neural subspaces supported by different coordination patterns across neurons). Yet, we also found shared neural dimensions between the cue and reward epochs, where the choice-direction information was consistently represented throughout the epochs. Those dimensions depended on the coherent temporal patterns of the single-neuron responses. These findings indicated that the temporally structured single-neuron responses worked as a dynamic population structure, supporting coherent representations throughout epochs of a given task.

It is recently suggested that choice-related encodings observed in higher-order cortical areas are not necessarily abstract signals but rather parsimoniously explained by combination of fundamental behavioral and contextual variables such as a specific spatial position and head angle^36^. Additionally, a recent study in rats revealed that neurons in the lateral entorhinal cortex are sensitive to subtle changes of egocentric view^37^. In contrast to these results, our analysis revealed that the choice directions were a better predictor for the individual PRC neural responses than other major variables previously noted, namely, body posture, non-orienting movements and spatial view (Supplementary Fig. 2). Although those individual factors could influence on the PRC responses, they were not unique to evoke the PRC responses. Thus, we conclude that the choice-direction encodings in the PRC is not derived from these single behavioral variables but rather an abstract signal integrating various computations related to the choice at a given epoch.

How does the nature of the choice-direction encodings differ between the cue and reward epochs? We further characterized those encodings by comparing between the correct and erroneous trials for each epoch (Fig. 4e– f). In the cue epoch, we did not find significant difference between choice-direction encodings in the correct and erroneous trials. Given that those neural responses are better explained by the choice directions than by head angles (Supplementary Fig. 2d), the cue-epoch responses might reflect internal signals such as decision or motor preparation driven by reward expectation^16, 47–50^. Contrary, in the reward epoch, the choice-direction encodings significantly decreased in erroneous trials. The reduction suggest that the presence of reward was necessary to elicit those responses.

Furthermore, the reward-epoch responses were temporally locked to the onset of reward and were not elicited by spatial position, view angle, reward expectation or combinations of them (Supplementary Fig. 2d–e). Therefore, those responses might convey reward signals in conjunction with choice directions. Those distinct choice-related representations were not expected by previous studies, most of which highlights perceptual and mnemonic representations in the PRC^18–22, 51–54^. However, a recent lesion study suggests that the PRC is also involved in value-based decision making^55^. In light of widespread anatomical connections of the PRC^23–25^, it is possible that this region carries diverse information besides perceptual and mnemonic signals depending on neural computation required for the task^56^. The above results emphasize different neural computations reflected in choice-direction selectivity in each of the cue and reward epochs. In spite of the explicit difference, the PRC neurons often showed choice-direction encodings in both epochs, suggesting that they support associative representations of choice and its outcome. This agrees with a traditional view, which holds that the PRC supports associative memory^14, 18–22^. Several studies showed that temporally sustained responses were prevalent in the PRC^16, 22–27^. Such sustained responses were found in studies focusing on neural responses during cue and memory epochs and were considered to serve as stable representations of targeted information across time epochs, which enable typical functions of the PRC including recognition memory^57^. Our finding of the inverted response patterns was not predicted by those previous studies but suggested its advantage in supporting both temporal coherence and flexibility of neural representations. By using the dimensionality reduction technique, we showed that the inverted encoding patterns of the PRC population mediated representations of the different epochs as well as consistent discriminability of the different choice-directions (Fig. 5c, g). Moreover, an important finding in our study is that the discriminability of the choice directions and task epochs was highly correlated (Fig. 5h). These results not only indicate dynamics of the PRC responses, which was underestimated in previous studies, but suggest an encoding strategy in the PRC efficiently enhanced discriminability of information from multiple sources across task epochs.

It is worth mentioning that values of neural weights of the individual neurons varied between the cue and reward epochs. In spite of the heterogeneity in individual neurons, at the population-level, the neural weights were moderately correlated between the epochs, resulting in the coherent choice-direction representations (Fig. 5e–f). Heterogeneous response properties across individual neurons are often observed in high-order cortical areas^29, 31, 38, 58–60^ and are considered to increase the number of dimensions can be represented by a neural population^8, 61–64^. In some cases, neurons exhibit a high degree of heterogeneity, in which multiple variables were randomly mixed at the individual neurons^6–7^. This ‘random mixed selectivity’ provide orthogonal neural subspaces where different neural computations can be performed independently. In light of these ideas, the moderately correlated neural subspaces between the cue and reward epochs exhibit balanced coherence and dynamics of the neural representations in the PRC, which might enable integrated processing of a targeted variable and contextual information as reported by previous studies^26–27^.

In conclusion, the PRC maintained choice-direction information while representing different epochs of a task. These representations were achieved via neural dimensions shared across the cue and reward epochs despite the dynamic reorganization of the population encoding structure. The neurons with coherent responses showed their heterogeneous contribution to those dimensions, indicating reconciled coherence and dynamics of neural representations across different levels.

## Methods

### Subjects

Seven male Long-Evans rats (Shimizu Laboratory Supplies, Kyoto, Japan) weighting 278–375 g at the beginning of the training were individually housed and maintained on a laboratory light/dark cycle (lights on 8:00 A.M. to 9:00 P.M.). Rats were placed on water restriction with *ad libitum* access to food. The animals were maintained at 80% of their baseline weight throughout the experiments. All experiments were implemented in accordance with the guidelines for the care and use of laboratory animals provided by the Animal Research Committee of the Doshisha University.

### Behavioral apparatus

The behavioral apparatus (Fig. 1a) has been previously described^13, 65^. An operant chamber (O’Hara, Tokyo, Japan) with three ports in the front wall for nose-poke responses was enclosed in a soundproof box (Brain Science Idea, Osaka, Japan). Each port was equipped with an infrared sensor to detect the animals’ nose-poke responses. Visual cues were presented using white light-emitting diodes (LEDs) (4000 mcd; RS Components, Yokohama, Japan) placed on the left and right walls of the operant chamber, as in Fig. 1a. Cue odors were presented via the central port through a stainless tube. The odors were mixed with pure air to produce a 1:10 dilution at a flow rate of 60 ml/min using a custom-built olfactometer (AALBORG, Orangeburg, NY). Water rewards were delivered from gravity-fed reservoirs regulated by solenoid valves (The Lee Company, Westbrook, CT) through stainless tubes placed inside of the left and right target-ports. We controlled stimulus and reward deliveries and measured behavioral responses using Bpod and Pulse Pal^66^ (Sanworks, Stony Brook, NY).

### Two-alternative forced-choice task

Each trial started when the rats poked their snout into the central port (Fig. 1b). After a variable delay (200–600ms, uniform distribution), a cue randomly selected from four stimuli (left/right LED for visual modality, S(+)/R(−)-2-octanol for olfactory modality) was delivered. If the rats successfully maintained their nose in the central port during 1 s after the cue onset, the “go” sound was delivered, and they were allowed to withdraw from the central port and to choose either left or right target port based on the task rule (bottom right in Fig. 1a). The presentations of the cue and the go sound were terminated by the withdrawal from the central port. When the rats left the central port without waiting for the go sound, the trial was canceled and followed by a 5 s punish intertrial-interval. Only correct choices were immediately rewarded by a drop of water (13 µl for five rats and 16 µl for two rats), from the target port. Rats performed 861 ± 232 trials in a daily recording session (seven rats, 50 sessions).

For two of seven rats (Supplementary Fig. 2), a 500 ms delay period was inserted between the onset of the target choice and the reward to assess the potential influence of non-orienting movements (licking) and spatial view on neural responses.

### Training

We trained the rats step-by-step to perform the task described above. The training period typically lasted 4 to 8 weeks. First, rats were trained to poke into the central port and then collect the water reward (20 µl) from the left or right target-port. We gradually extended the duration of the central poke by delaying the go sound up to 1 s after the poke onset. Next, the rats were trained to discriminate the odor cues based on the same contingencies as the recording sessions. A variable delay (200–600 ms) was inserted before the cue onset. After the rats became able to successfully discriminate the odor cues (> 80%), they were also trained to discriminate the visual cues based on the same contingencies as the recording sessions (> 80%). Finally, we interleaved visual and olfactory trials within a session and trained the animals to perform the task according to a training performance criterion (> 80%).

For the two rats (Supplementary Fig. 2), a reward delay period was introduced after they acquired the odor discrimination. The reward delay was gradually extended from 100 to 500 ms.

### Mixture of odors

The cue odors, S(+)/R(−)-2-octanol, were mixed together in a subset of sessions in order to increase the difficulty of the olfactory discrimination and thereby obtaining a sufficient number of erroneous trials. For instance, we used a 60/40 ratio in a given session, delivering an odor mixture of 60% S(+)-2-octanol and 40% R(−)-2-octanol. We maintained the odor discrimination accuracy constant (> 80%) throughout the recording sessions by adjusting the degree of odor mixing before each session.

### Surgery

Rats were anesthetized with 2.5% isoflurane before surgery, and it was maintained throughout surgical procedures. We monitored body temperature, movements and hind leg reflex and adjusted the depth of the anesthesia as needed. An eye ointment was used to keep the eyes moistened throughout the surgery. Subcutaneous scalp injection of a lidocaine 1% solution provided local anesthesia before the incision. The left temporalis muscle was retracted to expose the skull during the surgery. A craniotomy was performed over the anterior part of the left PRC (AP −3.5 to −3.24 mm, ML 6.6 to 6.8 mm relative to the bregma, 3.5 to 4.0 mm below the brain surface) and a custom-designed electrode was vertically implanted using a stereotactic manipulator. A stainless-steel screw was placed over the cerebellum and served as the ground during the recordings. We used the mean response of the all electrodes as a reference. During a week of postsurgical recovery, we gradually lowered the tetrodes to detect unit activities in the PRC. Electrode placement was estimated based on the depth and was histologically confirmed at the end of the experiments.

### Histology

Once the experiments were completed, the rats were deeply anesthetized with sodium pentobarbital and then transcardially perfused with phosphate-buffered saline and 4% paraformaldehyde. The brains were removed and post-fixed in 4% paraformaldehyde, and 100 μm coronal sections of the brains were prepared to confirm the recording sites.

### Electrophysiological recordings

A custom-designed electrode composed of eight tetrodes (tungsten wire, 12.5 µm, California Fine Wire, Grover Beach, CA) was used for the extracellular recordings. The tetrodes were individually covered by a polyimide tube (A-M Systems, Sequim, WA), were placed at a 100 µm separation and typically had an impedance of 150–700 kΩ at 1 kHz. The signals were recorded with Open Ephys (Cambridge, MA) at a sampling rate of 30 kHz and bandpass filtered between 0.6 and 6 kHz. The tetrodes were lowered approximately 80µm after each recording session, and thereby independent populations of neurons were recorded across the sessions.

### Monitoring and processing of body posture during task performance

We used a head-mounted accelerometer (Intan Technologies, Los Angeles, CA) to obtain postural signals from the animals (*n* = 2) during the electrophysiological recordings (Supplementary Fig. 2). The accelerometer signals (x-, y-, z-axis) were recorded at a sampling rate of 30 kHz and then downsampled to 100Hz. To precisely detect the body posture of the animals, gravity components of the accelerometer signals were estimated by using a low-pass filter with a cut-off frequency of 2 Hz^40^. To reduce the influence from the head angle that each animal typically preferred^40^, for each axis, the above processed signals were normalized to the mean and standard deviation in the baseline epoch (−400 to 0 ms before the cue onset).

### Spike sorting and screening criteria of units

All analyses were performed using MATLAB (MathWorks, Natick, MA). To detect single-neuron responses, the spikes were manually clustered with MClust (A.D. Redish) for MATLAB. Only neurons met the following criteria were included for further analyses: (1) units with sufficient isolation quality (isolation distance ≧ 15); (2) units with reliable refractory periods (violations were less than 1% of all spikes); and (3) units with sufficient mean firing rates in the 1 s after the cue onset (> 0.5 Hz). On average, we detected 10.9±7.4 neurons in a single recording session, and 6.2±3.4 neurons survived these quality criteria (total 312 neurons from seven rats).

### Selective responses to choice directions

In order to evaluate the selective responses to different direction of choices, we computed a choice-direction selectivity^67^. We first grouped correct trials into four types based on the cue modality (visual or olfactory) and target choice (left or right). For each modality, we independently calculated the choice-direction selectivity by using ROC analysis^68^. The choice-direction selectivity was obtained from the area under ROC curve (AUC) and defined as 2 × (AUC−0.5) ranging from −1 to 1. In our analysis, a positive value indicated a neuron selectively fired to the left target-choice, and a negative value indicated the opposite. A value of zero indicated the absence of choice-direction selective responses. To determine statistical significance (*P* < 0.05), we used permutation tests (1,000 interactions). For visualization of the temporal patterns of the choice-direction selectivity in individual neurons (Fig. 2a, Fig.3, Supplementary Fig. 1, and Supplementary Fig. 2b), mean firing rates were computed in 10 ms time windows (smoothed with a Gaussian, σ = 30 ms), and we then computed the choice-direction selectivity at each time point from those data.

### Statistics

We evaluated the statistical significance of the decoding analysis and the state space analysis with a bootstrapping procedure^8^. We estimated the *P* value for the bootstrapping procedure by computing the ratio (1+ *X*) / (*N* +1), where the number *X* indicates overlapping data points between the two distributions and the number *N* indicates interactions. Since we used 1,000 bootstraps, two distributions with no overlap resulted in *P* < 0.001, and two distributions with *x*% overlap resulted in *P* ≈ *x* /100. All the data are presented as mean ± standard deviation, unless otherwise stated.

### Decoding analysis

We employed a cross-temporal pattern analysis^42, 59, 69^ to investigate the temporal changes in the choice-direction selective responses in the PRC. Neural responses were pooled across the recording sessions to maximize the number of neural responses included in the decoding analysis. Here, we refer to the pooled pseudo-population of the PRC neurons (*n* = 207 from five rats) as the full population. The instantaneous firing rate of each neuron was estimated by the spike counts in a 150 ms sliding window (10 ms increment). We computed the above described choice-direction selectivity from the instantaneous firing rates for each neuron independently for visual and olfactory trials. In this manner, we generated two independent population vectors for the full population (cells × time matrix each for the cue modalities). We obtained a pattern similarity index by calculating the Fisher-transformed Pearson correlation (*r’*) between these two population vectors. This index provided the pattern similarity for both equivalent and different time points (for example, Fig. 4a). A positive value was interpreted as evidence for a reliable choice-direction encoding irrespective of the cue modality.

To estimate the mean performance values for the pattern classification analysis, we randomly resampled the neurons (the same number of neurons as neural population analyzed) and computed the choice-direction selectivity in visual and olfactory trials. Neural responses were aligned to withdrawal onset from the central port (for a cue epoch), target-choice onset (for a reward epoch) and target-choice offset (for two return-epochs). For return-epoch responses, we analyzed neural responses before and after the animals left the target ports in preparation for the next trials (pre-return epoch: −400 to 0 ms before leaving a target port: post-return epoch: 0 to 400 ms after leaving a target port). Only trials where the animals directly returned from the target port to the central port for the next trial were included in the analysis. These data allowed us to compute the pattern similarity indices within an epoch or across two different epochs. To investigate choice-direction encoding during the epochs, we averaged the performance within each of the epochs (the cue, reward, pre-return and post-return epochs). To obtain a baseline performance, we averaged the classification performance during the baseline epoch, −400 to 0 ms before the cue onset. We also quantified the pattern similarity of the neural responses between two epochs (for simplicity, we here refer to these epochs as A-epoch and B-epoch) by averaging the following two pattern classifications: pattern classification for A-epoch responses in visual trials and B-epoch responses in olfactory trials, and A-epoch responses in olfactory trials and B-epoch responses in visual trials (for example, Fig. 4c). We repeated the above processes 1,000 times to obtain a distribution of 1,000 different measurements of each pattern classification. To determine statistical significance, we compared a distribution obtained from 1,000 different pattern classification measurements within an epoch with a baseline distribution using the above described estimated *P* value. We considered zero to be chance level instead of the baseline distribution when we verified statistical significance of the classification across the epochs.

### Decoding analysis of erroneous trials

We included 119 neurons recorded in sessions with a sufficient number of erroneous target-choices (at least 11 trials for both directions of target-choice) in all the analyses for erroneous trials. Due to the similarity of the choice-direction selectivity between the different cue-modalities in correct trials (Fig. 2c and 4a), we computed the pattern similarity index using correct visual trials and erroneous olfactory trials to evaluate the influence of erroneous behavioral performance on the choice-direction encoding (Fig. 4e). In the erroneous trials, the animals chose the same target-port as the correct trials. To determine statistical significance of pattern classification performance in correct and erroneous trials, we downsampled the correct olfactory trials to match the number of trials for the erroneous olfactory trials (Fig. 4f). To test whether the classification performance during each epoch decreased from the correct trials to erroneous trials, we subtracted the classification performance in the erroneous trials from that in the correct trials for each resampling. The distribution of the residual performance was compared with zero by using the estimated *P* value.

### State space analysis

To understand the population structure and its temporal change between the cue and reward epochs, we performed principal component analysis (PCA). For each of the epochs, we constructed a 207 neurons × 4 conditions matrix^43–44^, in which columns contained trial-averaged z-scored firing rates of each neuron. The instantaneous firing rate of each neuron (estimated by spike counts in a 150 ms sliding window with 10 ms increment) was converted to a z-score by normalizing to the mean and standard deviation of its own instantaneous firing rates during the epoch. We then obtained time-averaged firing rates of the neurons each for the cue and reward epochs. By performing PCA on these datasets, we reduced the dimensionality of the PRC population from 207 neurons to two principal components (Supplementary Fig. 5a). We independently performed this analysis for population responses during the cue and reward epochs to obtain the cue-epoch and reward-epoch subspaces.

For data projections onto the above two-dimensional neural subspaces, we randomly selected 25 trials for each of the four conditions. z-scored firing rates of each neuron were then obtained by normalizing the data to the mean and standard deviation of its own firing rates during the epoch corresponding to the neural subspaces in which the projections were performed. We averaged the z-scored firing rates of each neuron during each of three epochs (the cue, reward and baseline epochs) to obtain a 207 cells × 4 conditions matrix for each epoch. To visualize the population responses, we projected these data onto the two-dimensional PCA space. This allowed us to obtain a single point reflecting the entire population response for each of the four conditions. We repeated this procedure five times with different subsets of 25 trials, and this allowed us to reduce some degree of the variability among individual trials. To obtain the within-condition distance, we computed the mean of pairwise Euclidean distances for five points in each of the four conditions and then averaged those distances across the conditions to obtain a single value for the within-condition distance. To obtain the across-condition distance, we computed pairwise distances between two sets of five points obtained from two different conditions in the same subspace and then averaged them. This procedure was repeated for all possible combinations of two different conditions, and we then averaged those distances to obtain a single value for the across-condition distance. We also investigated how the representational geometry changed between different epochs (that is, temporal shift of population responses in a neural subspace). This was achieved by computing the mean of pairwise distances between two sets of five points obtained from neural responses under a single condition in two different epochs. The mean distance was obtained for all trial conditions and then averaged to produce a single value for the across-condition distance for the temporal shifts.

To visualize variability across different subsets of trials (Fig. 5a) and test statistical significance, the above analysis was repeated 1,000 times with different subsets of resampled trials. The across-condition distances of the projections on corresponding subspaces (that is, cue-epoch responses onto cue-epoch subspace and reward-epoch responses onto reward-epoch subspace; Fig.5a) were compared with a distribution of the within-condition distance using the above described estimated *P* value (Fig. 5b). Across-condition distances larger than within-condition distance indicated that different trial conditions were discriminated at the population level.

Similarly, the across-condition distances of projections onto the interchanged subspaces (that is, cue-epoch responses onto reward-epoch subspace and reward-epoch responses onto cue-epoch subspace; Fig. 5a) were compared with a distribution of within-condition distance by using the estimated *P* value (Fig. 5c). For the projections onto the interchanged subspaces, we also projected baseline-epoch responses as negative controls (Fig. 5c and Supplementary Fig. 5c). To investigate the importance of the temporal response patterns of individual neurons, we projected randomly shuffled data (Fig. 5c and g), where the correspondence between the cue-epoch responses and reward-epoch responses were shuffled among the neurons (Supplementary Fig. 5d). We considered across-condition distances of the baseline-epoch responses to be chance when we evaluated the temporal shifts in representational geometry between the cue and reward epoch in a neural subspace (Fig. 5g).

To directly evaluate contribution of individual neurons to the choice-direction encodings (Fig. 5f), we performed the following analysis. We first sorted all the neurons from ones with the highest magnitude of the weighting values on the first dimension of the cue or reward epoch. For these differently sorted populations, we performed cluster projections onto the corresponding subspaces with increasing numbers of neurons. For precise evaluation of each neuron’s contribution, rather than performing PCA on each of differently sized populations independently, we used the same projection data in Fig. 5a (that is, the full population data) and replaced the weighting values of excluded neurons with zero. In each size of population, across-condition distance for different choice-directions on the first dimension was computed for each modality and then averaged. Those distances were then normalized to the distance obtained from the full population. We compared the distances from differently sized populations with those from the full population by the estimated *P* value. When the estimated *P* exceeded 0.05, we considered that number of neurons was sufficient to achieve discriminability of the choice directions equivalent to the full population.

Relationship between choice-direction and task-epoch discriminability To understand how the PRC neural population reconciled coherent representations of the choice directions (Fig. 5c) with dynamic changes for different task-epochs (Fig. 5g), we performed the following analyses. First, to quantify the degree of representational coherence in the PRC neural responses between the cue and reward epochs, the 1,000 across-condition distances of interchanged projections (for example, the reward-epoch responses onto the cue-epoch subspace shown in Fig. 5c) were divided by the 1,000 across-condition distances of the corresponding projections (for example, the cue-epoch responses onto the cue-epoch subspace shown in Fig. 5b). We repeated this procedure for projections of each of the cue and reward subspaces and then averaged between them to obtain the 1,000 normalized distances. Those normalized distances reflected the degree of discriminability of the different trial conditions preserved across the different epochs, and we referred those values as a temporal coherence index (x-axis in Fig. 5h). As negative controls, for each of the cue and reward subspace, the across-condition distances from the interchanged projections of the baseline-epoch responses and the randomly shuffled responses (Fig. 5c) were also divided by the across-condition distances of the corresponding projections (Fig. 5b), and we then averaged them between the epochs (Fig. 5h). Second, to quantify the degree of dynamic changes in the PRC neural responses between the cue and reward epochs, the 1,000 cluster distances obtained from temporal shifts of the population responses in a neural subspace (for example, distances from the cue-epoch responses to the reward-epoch responses in the cue-epoch subspace, which was computed for Fig. 5g) were divided by the 1,000 cluster distances obtained from simulated maximum temporal shifts of population responses in the same neural subspace (in this case, distances from the cue-epoch responses to the completely inverted cue-epoch responses in the cue-epoch subspace). The simulated neural responses were produced by inverting signs of the original projection data in the first and second dimensions. We repeated this procedure for projection data on each of the cue and reward subspaces and then averaged between them to obtain the 1,000 normalized distances.

Those normalized distances reflected the degree of discriminability of the different epochs (that is, the cue and reward epochs), we therefore referred those values as a temporal shift index (y-axis in Fig. 5h). Given the procedures for identifying neural subspaces by PCA, in which the neural responses were mean-centered in each epoch, we did not suppose any types of representational geometry that were not centralized or had temporal changes with deviation from the range of across-condition distances in the corresponding projections. Therefore, the completely inverted response pattens provided the maximum level of temporal shifts (temporal shift index = 1). This also indicates that the temporal shift index equals 0.5 when projected clusters overlap on the center of neural subspaces. As negative controls, for each of the cue and reward subspace, the temporal shift distances from the interchanged projections of the baseline-epoch responses and randomly shuffled responses (Fig. 5g) were also divided by the cluster distances obtained from the above described simulated temporal shifts of population responses and then averaged between the epochs.

## Data availability

The data sets analyzed in the present study are available from the corresponding authors upon reasonable request.

## Code availability

The code used for analysis in the present study are available from the corresponding authors upon reasonable request.

## Acknowledgments

We would like to thank members of the Laboratory of Neural Information at Doshisha University for helpful discussion, and Y. Tanisumi for help with visualization of the data. This research was supported by the JSPS KAKENHI Grant Numbers 19J12634 (to T.O.), 16H02061, 18H05088 (to Y.S.) and 16K18380 (to J.H.).

## Author contributions

T.O., Y.S., and J.H. designed the experiments. T.O. performed the experiments and analyzed the data. Y.O. analyzed the data. H.M, Y.S. and J.H. supervised the project. All authors contributed to writing the manuscript.

## Competing interests

The authors declare no competing interests.

