## Supplemental Data 1 for "Dynamic coordination of the perirhinal cortical neurons supports coherent representations between task epochs"

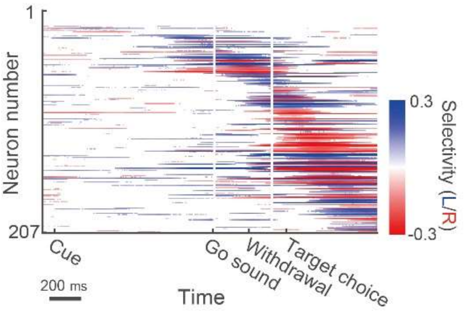


**Supplementary Figure 1 | Choice-direction encodings in olfactiry trials.** Temporal patterns of choice-direction selective responses of the PRC neurons ($n$ = 207) in olfactory trials. Selective responses to left choice were shown in blue, and the opposite were shown in red. Neural responses in correct trials were shown. Only segments with significant selectivity were shown ($P$< 0.05; 1,000 permutations).

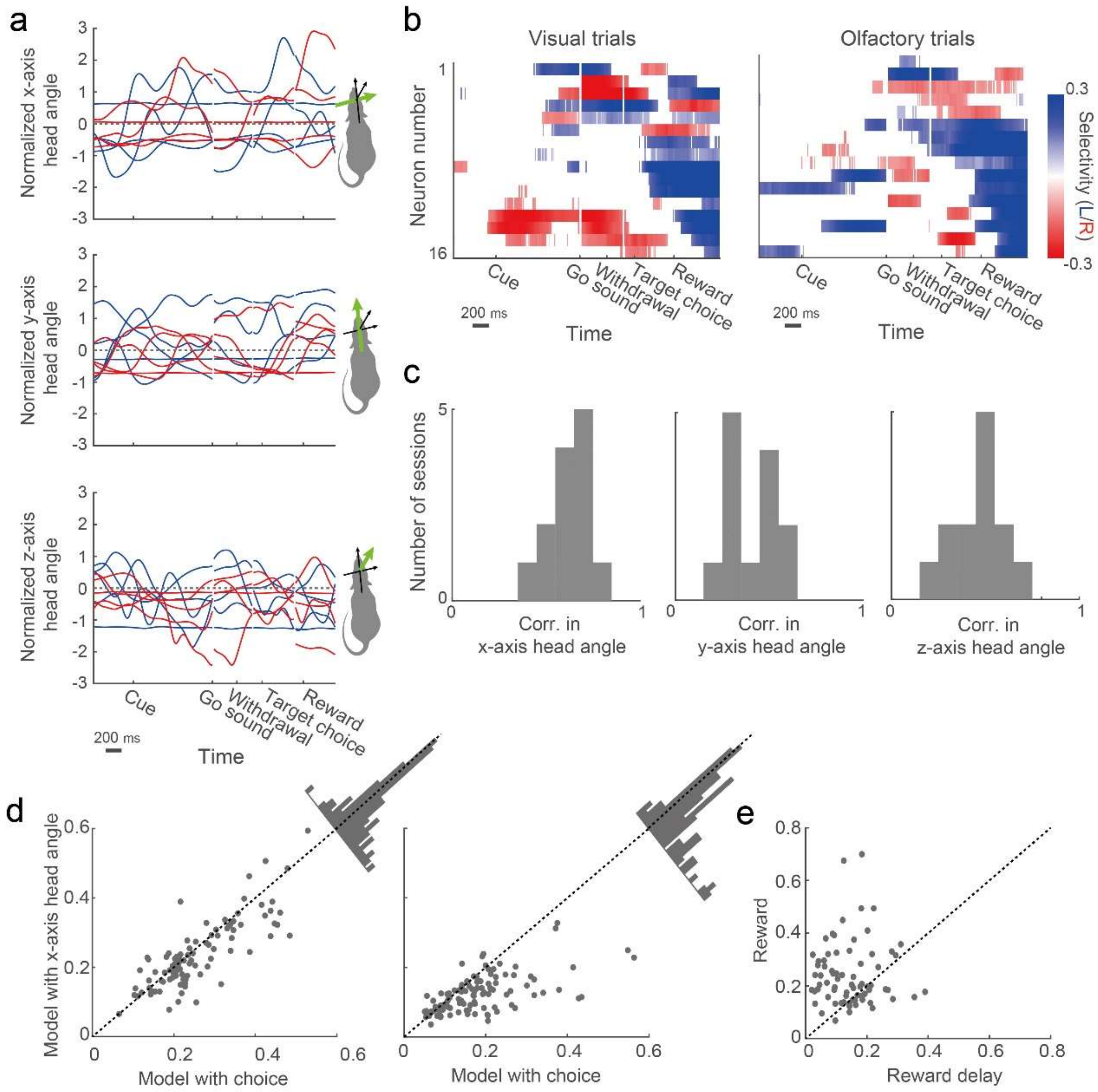


**Supplementary Figure 2 | Influence of fundamenral behavioral and contextual factors on choice-direction encodings in the PRC.** (a) Diverse patterns of head angles of a rat in left (blue) and right (red) trials of an exapmle session (463 trials). Traces represent estimated gravity components of accerelometor signals (Methods) in 5 trials randomly sampled for each direction of the target choice. (b) Choice-direction selectivity of neurons ($n$ =

16) recorded in the example session shown in a. Only segments with significant selectivity are shown ($P < 0.05$; 1,000 permutations). For each modality, neurons were sorted according to their peak selectivity. In this representative session, seven neurons inverted the choice-direction selectivity between the cue and reward epochs. Note that the reward epoch was defined as 0 to 400 ms after the onset of reward. (c) Correlation of head angles between the cue and reward epochs always showed positive values, suggesting no temporal inversion of body posture (13 sessions from two rats). (d) Comparison of prediction performance between linear-regression models with choice direction and x-axis head angle ($n$ = 105 neurons from two rats) in the cue epoch (left) and the reward epoch (right). Those models also included y-axis head angle, z-axis head angle, and reaction time as expranatory variables. For each neuron (shown by a point), firing rate during an epoch was independently predicted by the two models. The performance of the models were evaluated by computing correlation coeffients between the neural responses and the model prediction across trials. The better model to explain the neural responses was determined by the difference in correlation values between the models (the distributions are shown by histograms). (e)

Comparison of the magnitude of the choice-direction selectivity between the reward epoch and the reward-delay epoch (0 to 400 ms after the target-choice onset). Neurons with significant choice-direction selectivity ($P < 0.05$; 1,000 permutations) during the reward epoch were included in this analysis ($n = 73$ of 105 neurons).

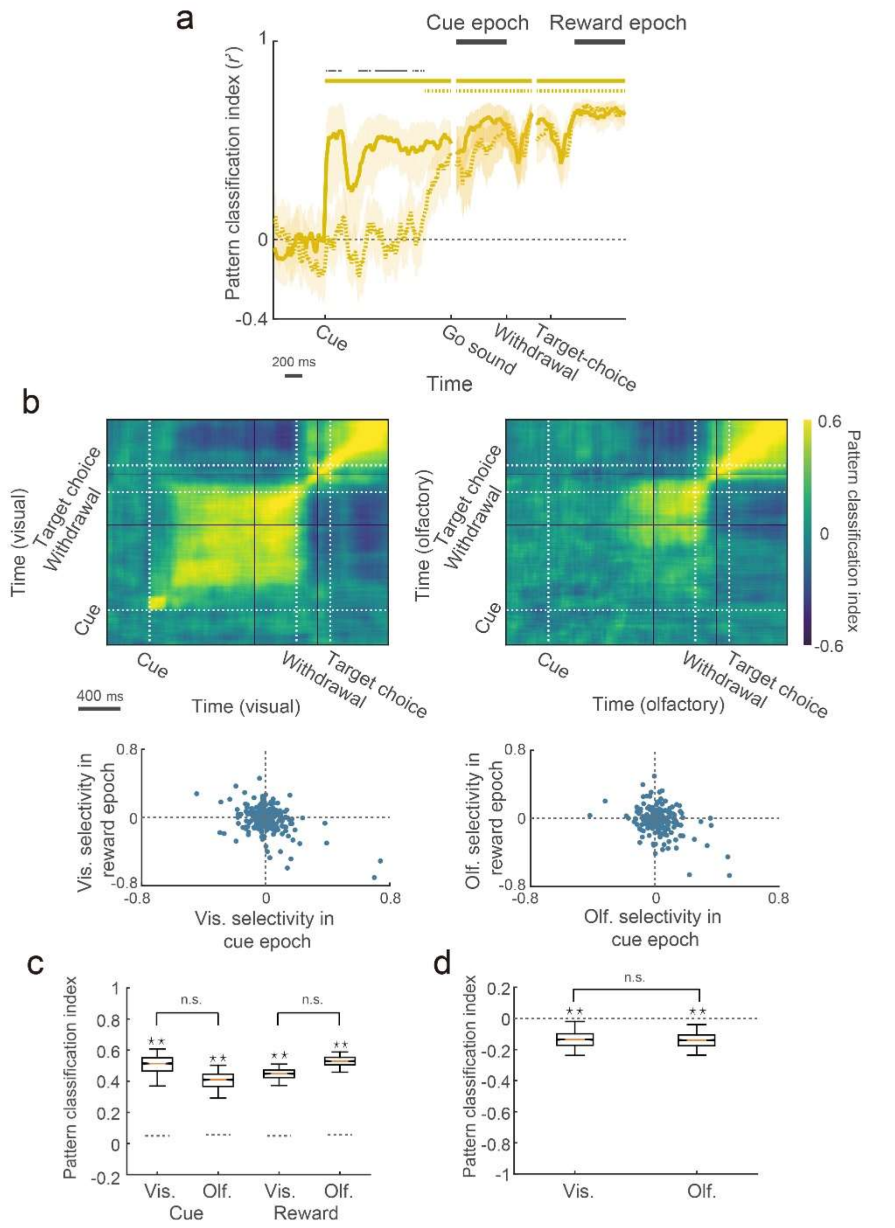


**Supplementary Figure 3 | Time-resolved pattern analysis in each cue-modality.** (a) Performance of the time-dependent pattern classification analysis obtained independently from visual and olfactory trials (shown by solid line and dashed line, respectively). In each modality, the classification performance for different choice-directions was computed on the neural data which was subdivided into two groups by an interleaved approach (odd trials for train data and even trials for test data). The middle yellow lines and the surrounding shaded areas indicate mean performance and the 95th percentile range (two-sided) of the performance, respectively. The upper yellow solid (visual) and dashed (olfactory) lines correspond to time points in which the classification performance in each modality was above chance. The statistical significance in each time point was determined by comparing the classification performance with zero (estimated $P < 0.05$; gray dashed line). The upper gray dots indicate time points where the classification performance was significantly different between the visual and olfactory trials (estimated $P < 0.05$). The classification performance increased with different latency between the modalities but reached to an equivalent level during the cue and reward epochs (thick black lines). (b) Cross-temporal pattern analysis for visual (upper left) and olfactory (upper right) trials. Temporal inversion of choice-direction selectivity between the cue and reward epochs are shown in bottom left (visual trials) and right (olfactory trials). (c) Mean classification performance during the cue and reward epochs in each modality. Dashed line indicates the 97.5th percentile values of the baseline epoch performance. In both epochs, the performance was significantly higher than the baseline performance ($P < 0.001$ for cue-epoch in visual trials; $P < 0.001$ for cue-epoch in olfactory trials; $P < 0.001$ for reward-epoch in visual trials; $P < 0.001$ for reward-epoch in olfactory trials) and was similar level between the modalities ($P \approx 0.995$ for cue-epoch; $P \approx 0.8931$ for reward-epoch). (d) Mean classification performance across the cue and reward epochs in each modality. In both modalities, the PRC showed reliable inversion of choice-direction encoding patterns ($P < 0.014$ for visual trials; $P < 0.003$ for olfactory trials). The inverted encoding patterns were similar level regardless of the modalities ($P \approx 0.998$). In box plots: orange line, median; box limits, 25th and 75th quartiles; notch limits, (1.57×interquartile range)/$\sqrt{n}$; whiskers, 95th percentile range (two-sided) of the distribution. Asterisks indicate statistical significance based on estimated $P$ values ($P < 0.05$; Methods), and n.s. indicate insignificance.

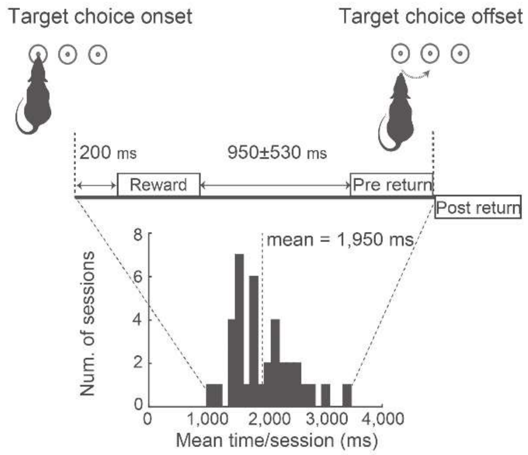


**Supplementary Figure 4 | Temporal relationships among the reward, pre-return, and post-return epochs.** The rats typically stayed in the target port 1950 ± 530 ms (mean ± s.d.) after the target-choice onset (37 sessions in five rats). The reward and pre-return epoch were separated by a time period ranging 950 ± 530 ms (mean ± s.d.).

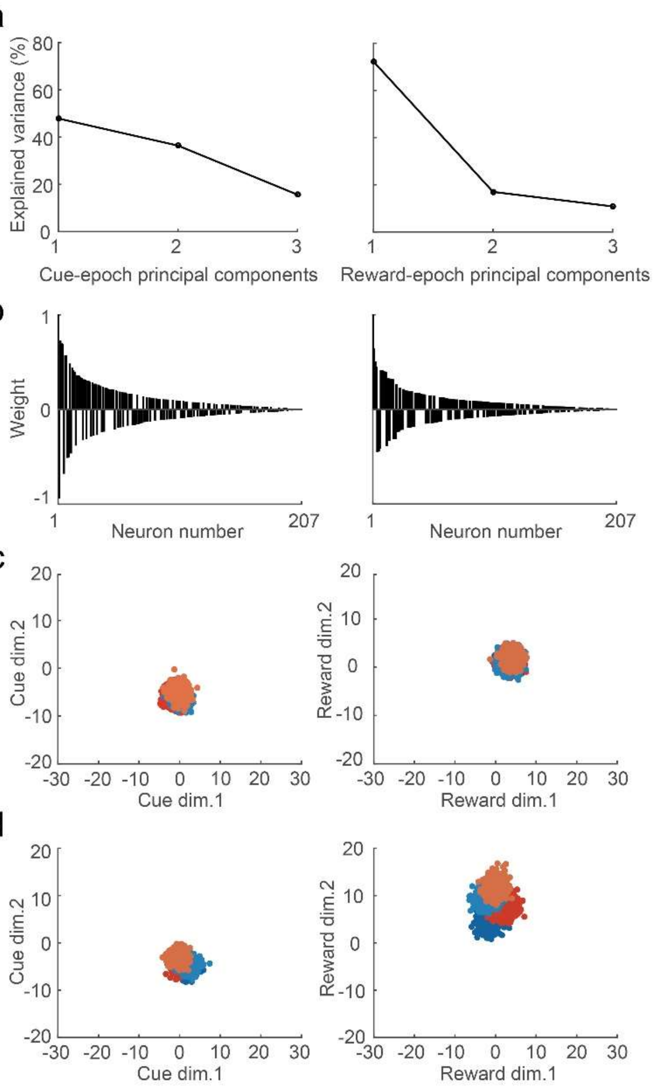


**Supplementary Figure 5 | Principal component analysis for population responses.** (a) Percentage of the cue-epoch response variance explained by the cue-epoch principal components (left) and percentage of the reward-epoch response variance explained by the reward-epoch principal components. The cue-epoch and reward epochs subspaces in Fig. 5 were defined by the first two PCs, which captured 84.36% and 89.18% of the data variance of the cue-epoch and reward-epoch responses, respectively. (b) Neural weights in second dimensions of the cue-epoch (left) and reward-epoch (right) responses. (c) Population responses during the baseline epoch (−400 to 0 ms before the cue onset) projected onto the cue-epoch and reward-epoch subspaces. Population responses in different conditions are shown in different colors: blue, left target-choice in visual trials; pale blue, left target-choice in olfactory trials; red, right target-choice in visual trials, orange, right target-choice in olfactory trials. (d) Projections of shuffled data onto interchanged subspaces. The shuffled data were generated by randomly shuffling order of neurons in the projection data shown in Fig. 5a. The shuffled reward-epoch responses were projected onto the cue-epoch subspace (left), and the shuffled cue-epoch responses were projected onto the reward-epoch subspace (right). Within-condition and across-condition distances obtained from c and d were shown in Fig. 5c and 5g.
